# Applying polygenic risk scoring for psychiatric disorders to a large family with Bipolar Disorder and Major Depressive Disorder

**DOI:** 10.1101/103713

**Authors:** Simone de Jong, Mateus Jose Abdalla Diniz, Andiara Calado Saloma Rodrigues, Ary Gadelha, Marcos L Santoro, Vanessa K Ota, Cristiano Noto, Major Depressive Disorder and Bipolar Disorder Working Groups of the Psychiatric Genomics Consortium, Charles Curtis, Hamel Patel, Lynsey S Hall, Paul F O’Reilly, Sintia I Belangero, Rodrigo Bressan, Gerome Breen

## Abstract

We aim to investigate the application of polygenic risk scoring within a family context. Polygenic risk profiles could aid in unraveling the role that common variation confers on disease risk within a pedigree that would have traditionally been viewed through the prism of monogenic inheritance only. We illustrate our discussion by analyzing polygenic risk scores for schizophrenia, major depressive disorder and bipolar disorder in a large pedigree (n~260) in which 30% of family members suffer from major depressive disorder or bipolar disorder. We apply polygenic risk scores to study patterns of assortative mating and anticipation, whereby it appears increased polygenic risk for psychiatric disorders is contributed by affected individuals who married into the family, resulting in an increasing genetic risk over generations in the family. This may explain the observation of anticipation in mood disorders, whereby onset is earlier and the severity of a disease increases over the generations of a family. Joint analyses of both rare and common variation may be the most powerful way to understand the familial genetics of mood and psychiatric disorders.

## INTRODUCTION

The development of polygenic risk scoring (PRS) has greatly advanced the field of psychiatry genetics. This approach allows for even sub-genome-wide significant threshold results from large genome wide meta analyses to be leveraged to explore genetic risk in smaller studies (Shaun M Purcell et al. 2009). The effect sizes at many individual single nucleotide polymorphisms (SNPs), estimated by a large genome wide association studies (GWAS) on the disorder of interest, are used to calculate an individual level genome-wide PRS in individuals from an independent genetic dataset. The PRS based on the summary statistics of the schizophrenia (SCZ) GWAS by the Psychiatric Genomics Consortium (PGC) (Ripke et al. 2014; Sullivan 2010) has proven to be most powerful in predicting not only SCZ (Ahn et al. 2016; S M Purcell et al. 2009) but also other psychiatric disorders (Lichtenstein et al. 2009; Lee et al. 2013; Cross-Disorder Group of the Psychiatric Genomics Consortium 2013). In addition, updated, more powerful, summary statistics from the Psychiatric Genomics Consortium from the latest GWAS for Bipolar Disorder (BPD) and Major Depressive Disorder (MDD) are available via the PGC Data Access Portal (https://www.med.unc.edu/pgc/shared-methods).

Aside from increasing power in traditional case-control designs, PRS algorithms also open up new avenues for studying common variation. In this study we consider the application of PRS within a family context. While pedigree studies have been traditionally used to explore rare genetic variation through linkage analyses, studying patterns of PRS throughout a pedigree would allow for assessment of phenomena like assortative mating and anticipation. Assortative (non-random) mating is a common phenomenon where mated pairs are more phenotypically similar for a given characteristic than would be expected by chance (Merikangas and Spiker 1982). Results from a recent study by Nordsletten et al. (Nordsletten et al. 2016) show extensive assortative mating within and across psychiatric, but not physical disorders. This could explain some of the features of the genetic architecture of this category of disorders (Nordsletten et al. 2016; Plomin, Krapohl, and O’Reilly 2016; Robinson et al. 2017). This includes anticipation, a phenomenon where later generations exhibit more severe symptoms at an earlier age, robustly reported (although not explained) in bipolar disorder (O’Donovan, Jones, and Craddock 2003) and recently highlighted in genetic studies of MDD (Power et al. 2016; Power et al. 2012).

In the current study we aim to discuss the application of polygenic risk scoring for SCZ, MDD and BPD to explore patterns of common risk variation within a family context. We illustrate our discussion by investigating the relationship between PRS and apparent assortative mating and anticipation within a complex multigenerational pedigree affected with mood disorders.

## MATERIALS & METHODS

### Subject description

The Brazilian Bipolar Family (BBF) was ascertained via a 45-year old female proband who presented with severe Bipolar Type 1 (BPI) disorder and stated there were dozens of cases of mood disorders in the family, most of whom lived in a small village in a rural area of a large state north of São Paulo. Cooperation from the family and a 2003 self-published book about their history was invaluable for our ascertainment. Historically, the entire BBF consists of 960 members. Living family members >16 years of age underwent semi-structured interviews, using the Portuguese version of the Structured Clinical Interview for DSM-IV Axis I Disorders (SCID-I) (Del-Ben, Rodrigues, and Zuardi 1996). Members aged 6-16 were assessed using the Portuguese version of Kiddie-SADS-Present and Lifetime Version (K-SADS-PL) (Brasil and Bordin 2010). In total 308 interviews were completed, and 5 eligible members declined an interview. In the rare event of discrepancies, two independent psychiatrists reviewed them and a final consensus diagnosis was assigned. All affected and unaffected adult family members that have been included in the genetic study have given informed consent. Minors have given assent, followed by consulted consent by their parents in accordance with accepted practice in both the U.K. and Brazil. The project was approved by the Brazilian National Ethics Committee (CONEP). Table 1 contains the demographics of the subjects used in the current analysis (n=243 passed genotype quality control procedures described below). The population control dataset (BRA controls) was collected in Sao Paulo, Brazil, as a control dataset in a genetic study of first-episode psychosis (Noto et al. 2015). They were volunteers who had no abnormal psychiatric diagnoses (SCID) or family history of psychotic illness. The Research Ethics Committee of Federal University of Sao Paulo (UNIFESP) approved the research protocol, and all participants gave informed consent (CEP No. 0603/10). Demographics for n=57 BRA controls can be found in Table 1.

**Table 1:** Demographics of the Brazilian Bipolar Family members and the Brazilian population control dataset (BRA controls) in the current study. Diagnostic categories are Bipolar I (BPI), Bipolar II (BPII), Bipolar not otherwise specified (BPNOS), recurrent Major Depressive Disorder (rMDD), Major Depressive Disorder (MDD), Schizoaffective disorder (SADB), Schizophrenia, Cyclothymia and Dysthymia. A breakdown of gender, age, age at onset is given in the next columns. The married-in column contains the number of individuals in each diagnostic category married-in to the family. The last column contains counts of individuals in each category who have experienced a psychotic episode during their lifetime.

| Diagnosis | n | Male, Female | Age (±sd) | Age of onset (±sd) | Married-in | Psychosis |
| --- | --- | --- | --- | --- | --- | --- |
| BPI | 17 | 6, 11 | 50.4 (±18.9) | 24.9 (±14.6) | 0 | 13 |
| BPII | 11 | 4, 7 | 38.7 (±15.2) | 24.2 (±13.8) | 1 | 4 |
| BPNOS | 8 | 6, 2 | 29.6 (±19.9) | 17.0 (±18.7) | 0 | 1 |
| rMDD | 17 | 5, 12 | 50.2 (±16.7) | 27.3 (±14.1) | 3 | 4 |
| MDD | 21 | 11, 10 | 43.8 (±17.8) | 34.5 (±15.5) | 6 | 1 |
| SADB | 1 | 0, 1 | 73 | 44 | 0 | 1 |
| Schizophrenia | 1 | 1, 0 | 44 | 36 | 0 | 1 |
| Cyclothymia | 1 | 0, 1 | 40 | 25 | 0 | 0 |
| Dysthymia | 1 | 0, 1 | 52 | - | 1 | 0 |
| Unaffected | 147 | 89, 58 | 36.8 (±20.0) | - | 35 | 0 |
| Unknown | 18 | 14, 4 | 5.7 (±7.1) | - | 0 | - |
| **Total** | **243** | **136, 107** | **37.3 (±21.0)** | **28.3(±15.5)** | **46** | **25** |
| BRA controls | 57 | 33, 24 | 27.1 (±7.2) | - | - | - |

### Genotype data

Following diagnostic interview, interviewers obtained whole blood in EDTA containing monovettes for adults and lesser amounts or saliva given personal preference or age (DNA Genotek Inc., Ontario, Canada). Genomic DNA was isolated from whole blood and saliva at UNIFESP using standard procedures. Whole genome genotype data was generated using the Illumina Infinium PsychArray-24 (http://www.illumina.com/products/psycharray.html) for both the BBF and the BRA control dataset at the in-house BRC BioResource Illumina core lab according to manufacturers protocol. Samples were excluded when average call rate was <98%, missingness >1% with additional check for excess heterozygosity, sex, family relationships and concordance rates with previous genotyping assays. SNPs were excluded when missingness >1%, MAF <0.01 or HWE <0.00001 and if showing Mendelian errors for the BBF dataset in Plink v1.07(S. Purcell et al. 2007) and v1.9 (Chang et al. 2015) or Merlin v1.1.2 (Abecasis et al. 2002). The BBF and BRA control datasets were QC’d separately and then merged, applying the same SNP QC thresholds to the merged dataset as well. This quality control procedure resulted in a dataset of 225,235 SNPs for 243 BBF individuals (197 family members and 46 married-in individuals) and 57 BRA controls. Eigensoft v4.2 (Patterson, Price, and Reich 2006) was used to check for population differences between the BBF family members, married-in individuals and BRA control sets. The BBF members self-reported mixed Southern European ancestry, confirmed by genome-wide principal components analysis showing that family members clustered closely with the Northern and Western European and Tuscan Italian populations in Hapmap3, with a relative lack of African or Native American ancestry (Supplementary Information S2). The principal components appear to represent within-family structure, with most PCs seemingly separating subfamilies (Supplementary Information S3 and S4). PC1 and PC2 are significantly correlated to the SCZ:PRS (PC1 *r*=-0.21, *p*=1.7 x 10^-4^; PC2 *r*=-0.22, *p*=1.4 x 10^-4^), MDD:PRS (PC1 *r*=0.12, *p*=0.04; PC2 *r*=-0.24, *p*=2.9 x 10^-5^) and BPD:PRS (PC1 *r*=0.19, *p*=8.8 x 10^-4^; PC2 *r*=-0.12, *p*=0.03). The principal components were not used in subsequent analyses.

### Polygenic risk scores

Polygenic risk scores for each family member (n=243) and population control (n=57) were generated in the same run using the PRSice v1.25 software (Euesden, Lewis, and O’Reilly 2014) with the publically available PGC schizophrenia GWAS (Ripke et al. 2014) as a base dataset (36,989 SCZ cases, 113,075 controls), in addition to MDD (51,865 MDD cases, 112,200 controls, not including 23andme individuals) and BPD (20,352 BPD cases, 31,358 controls) summary statistics from the latest PGC meta-analyses (unpublished data (PGC, Wray, and Sullivan 2017; Stahl et al. 2017)). We performed *P*-value-informed clumping on the genotype data with a cut-off of r^2^ = 0.25 within a 200-kb window, excluding the MHC region on chromosome 6 because of its complex linkage disequilibrium structure. Through linear regression (no covariates) in PRSice we selected the PRS thresholds most predictive in discriminating affected from unaffected family members for SCZ:PRS (*P<0.*00055, 1,218 SNPs), MDD:PRS (*P*<0.0165, 715 SNPs) and BPD:PRS (*P*<0.00005, 143 SNPs). PRS showed low to modest correlations (no covariates) amongst each other in our data (SCZ:PRS vs. MDD:PRS *r*=0.17, *p*=0.002, SCZ:PRS vs. BPD:PRS *r*=0.12, *p*=0.03, MDD:PRS vs. BPD:PRS *r*=-0.03, *p*=0.66).

### Linkage analysis

The main linkage analyses identifying rare genetic risk variation were performed as part of a previous paper on the BBF (Diniz et al. 2017) using the Affymetrix 10k linkage genotyping array. In order to explore the balance between common and rare risk variation we selected the strongest signal for affected versus unaffected family members on chr2p23 (chr2:30000001-36600000, LOD=3.83). Following the strategy described by Rioux et al. (Rioux et al. 2001) we performed a transmission disequilibrium (TDT) test on the 25 markers in this linkage region in an attempt identify ‘linkage positive’ individuals in n=300 family members with one or both types of genotype array data. N=155 individuals overlap with the current study and based on exploration of patterns of PRS in the current study we attempted to answer two questions: 1) with an increase of common risk variation, does rare risk variation become less important over generations, 2) do linkage positive individuals carrying the presumed risk allele show differences in PRS.

### Statistical testing

All PRS were standardized mean=0 and SD=1. Linear mixed model analyses were selected to be able to model covariates and relatedness within this complicated dataset. The analyses were performed using the Wald conditional F-test (Kenward and Roger 1997) in ASReml-R software (Butler et al. 2002) with one of the categories of mood disorders or family status as dependent variable and PRS as the independent variable. The model was adjusted for age (except for the generation analysis) and sex; and For 7 individuals in the BBF age at collection was missing and imputed to be the mean age of the relevant generation. To account for relatedness in within family comparisons, an additive genetic relationship matrix was fitted as a random effect. The relationship matrix was constructed using LDAK software (Speed et al. 2012) with weighted predictors and LD correction parameters suited for pedigree data, resulting in pairwise relatedness estimates and inbreeding coefficients on the diagonal. The variance explained by each PRS was calculated using: (var(x × β))/var(y), where x was the standardized PRS, β was the corresponding regression coefficient and y was the phenotype (Nakagawa and Schielzeth 2013). For the analysis of offspring of different spouse pair categories (“both unaffected”, “married-in parent affected”, “family parent affected”, “both affected”) we had to account for the number of children contributed to the same category by each spouse pair. While most spouse pairs contribute 1 or 2 children to the same offspring category (Supplementary Information S5); two “both affected” spouse pairs contribute 7 and 8 children respectively. In the event of more than one child per couple, we calculated the mean PRS per spouse pair and entered this in the model as being one representative child for that couple. All *p*-values reported are uncorrected for multiple testing, since all tests concern overlapping individuals and thus have a complex dependence structure. However, we have performed 42 tests as listed in Supplementary Information S6, and so a conservative Bonferroni threshold for *P* < 0.05 is 0.001.

## RESULTS

### Affection status

The PRS thresholds were selected to optimally discriminate between affected (n=78) versus unaffected (n=147) family members with a higher score in affecteds for SCZ:PRS (Beta=0.07, SE=0.03, Z-ratio=6.58, *p*=0.03, R^2^=0.03), MDD:PRS (Beta=0.07, SE=0.03, Z-ratio=6.58, *p*=0.03, R^2^=0.02) and BPD:PRS (Beta=0.09, SE=0.02, Z-ratio=6.81, *p*=0.002, R^2^=0.04). None of the PRS significantly discriminated between individuals having experienced a psychotic episode at some point in their lives (n=25) versus the unaffected group (n=147). Find visualization of PRS in different diagnostic categories in Supplementary Information S7.

### Assortative mating

Married-in individuals were defined as individuals married to a BBF member, but having no parents in the family themselves. Of the 70 married-in individuals ascertained (irrespective of having genotype data) 19 (27%) were affected with a psychiatric disorder. This is significantly higher than the 17% population prevalence of the most common of the three disorders: MDD (Fisher’s exact *p*=0.02) (Silva et al. 2014). The unaffected married-in group does not differ from the general healthy population as evidenced by no significant differences in PRS as compared to the BRA control group. The above led us to investigate whether we can observe assortative mating on a genetic level, using PRS. In spouse pairs, we were unable to predict the PRS of the husband, using that of his wife, even when selecting concordant (both affected or both unaffected) pairs only. We considered the possibility that the married-in individuals might confer a different genetic predisposition to mood disorders to their offspring than the original family members. Demographics of the offspring in the different offspring categories (no affected parents (n=54); one affected family member parent (n=69); one affected married-in parent (n=15) and two affected parents (n=38)) are given in Supplementary Information S5 and S8. Indeed, we find that offspring of an affected married-in parent show increased SCZ:PRS (Beta=0.21, SE=0.03, Z-ratio=4.42, *p*=0.002, R^2^=0.19, Figure 1) and BPD:PRS (Beta=0.17, SE=0.03, Z-ratio=4.42, *p*=0.01, R^2^=0.13, Figure 1) as compared to having no affected parents.

**Figure 1:**
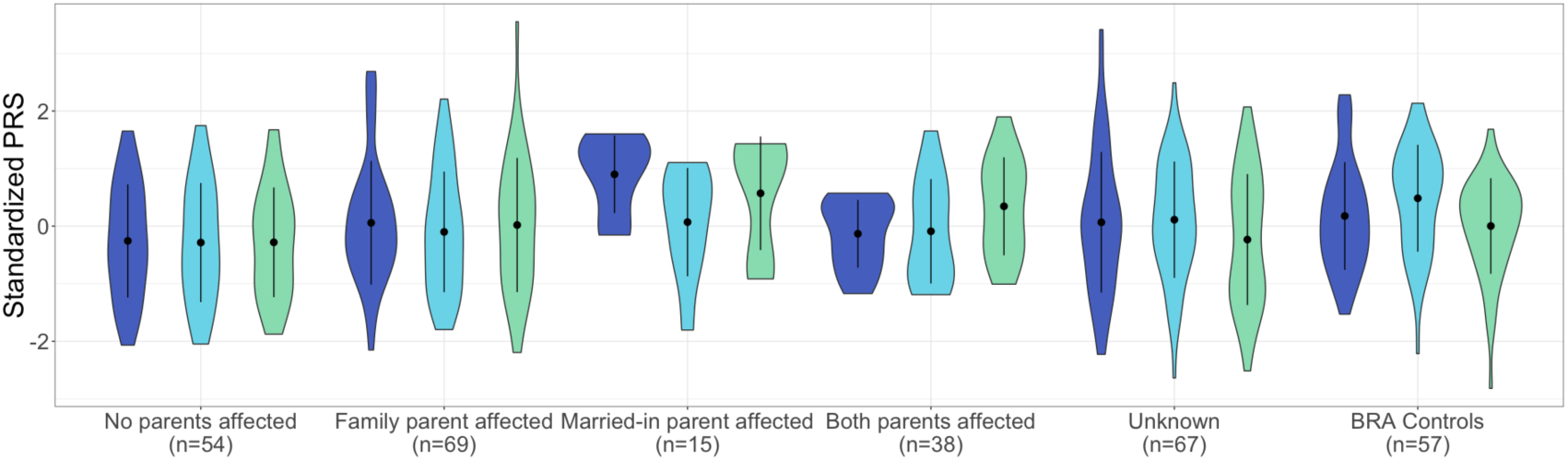
Violin plots of SCZ:PRS (dark blue plots) MDD:PRS (light blue plots) and BPD:PRS (green plots) for offspring of all spouse pair possibilities. The first category represent PRS in individuals with no affected parents, the next for individuals with an affected family member parent, followed by offspring of an affected married-in individual and finally offspring of two affected parents. The last two sets of violin plots represent offspring of unknown spouse pairs and the BRA controls. The dot and error bars represent mean +/- standard deviation of standardized PRSs.

### Anticipation

The BBF shows patterns of anticipation, with individuals having an earlier age at onset (AAO) in later generations. For 104 individuals (irrespective of having genotype data) the average age at onset significantly decreases over generations with G2 (n=1, AAO=8), G3 (n=23, AAO=30.2yrs±21.1), G4 (n=53, AAO=31.2yrs±12.3), G5 (n=23, AAO=19.7yrs±9.5) and G6 (n=4, AAO=13yrs±3.6) (Supplementary Information S9) with older participants recalling their AAO directly and younger participants confirmed using clinical records or parental recall (Beta=-4.55, SE=1.79, Z-ratio=-2.54, *p*=0.01, R^2^=0.06). We hypothesized that this decrease in AAO would be reflected in a negative correlation with PRS, subsequently resulting in a pattern of increased PRS over generations. Because of a limited sample size of affected individuals per generation, a direct correlation of AAO and PRS does not reach significance, although the youngest generation (G5) does show trends towards negative correlations for SCZ:PRS and MDD:PRS (Supplementary Information S10). The SCZ:PRS does show a significant increase over generations (Figure 2) where n=197 family members were included (46 married-in individuals were excluded from the analysis to capture inheritance patterns of SCZ:PRS) in a linear regression with generation as independent variable (Beta=0.13, SE=0.02, Z-ratio=0.85, *p*=0.008, R^2^=0.03). The presence of such an effect when comparing generations suggests ascertainment effects such as relying on the recall of older family member with very long duration of illness in previous generations may be masking an overall effect across the entire family.

**Figure 2:**
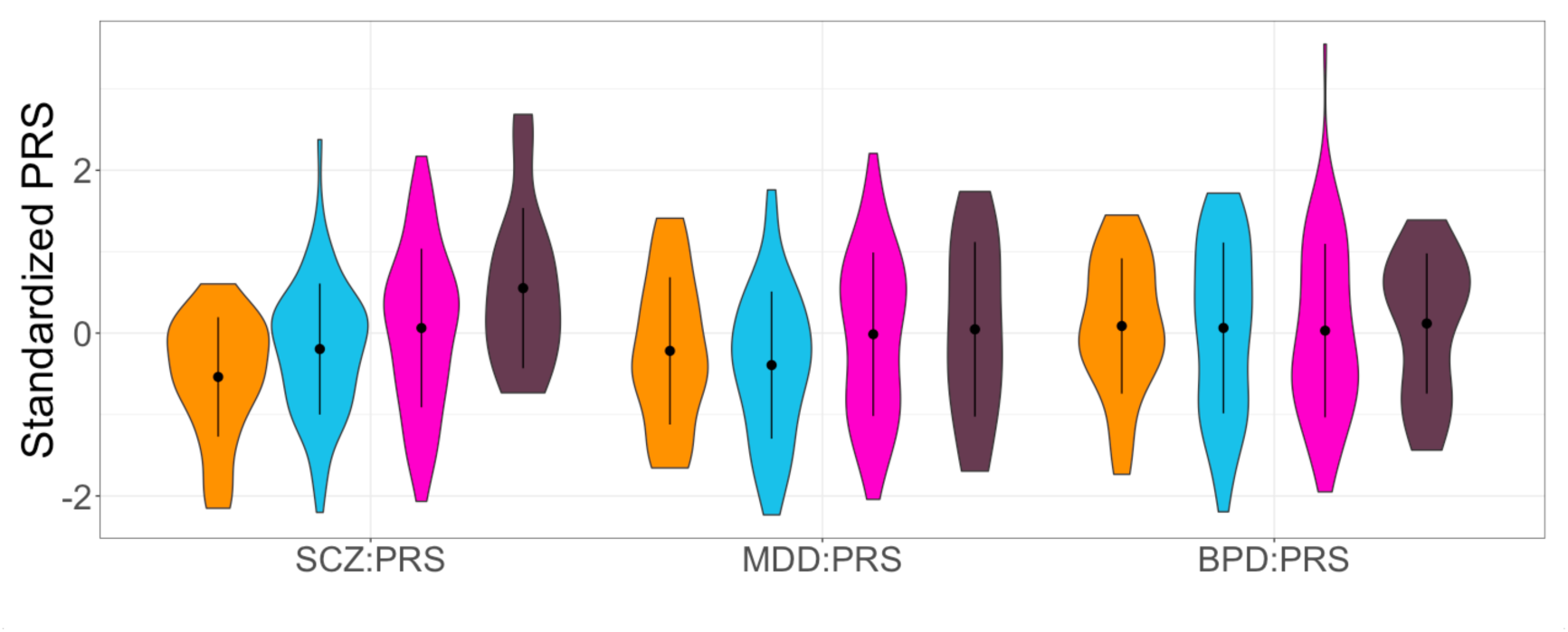
Violin plots of SCZ:PRS, MDD:PRS and BPD:PRS per generation for family members only, with results for the generations G3 (n=25, orange plots), G4 (n=72, light blue plots), G5 (n=80, pink plots) and G6 (n=16, dark purple plots) (excluding the oldest generation G2 and youngest generation G7 because of n=2 sample size). The dot and error bars represent mean +/- standard deviation of standardized PRSs.

### Balance of common and rare genetic risk

TDT analysis within the chr2p23 linkage region resulted in identification of rs1862975, a SNP originally typed on the Affymetrix linkage array (combined test *p*=0.003). The homozygous T genotype was detected in 68% affected family members, 57% affected married-ins, 36% unaffected family members and 24% unaffected married-ins. Since this SNP was present only on the Affymetrix SNP genotypes are not available for the BRA controls. The distribution of the rs1862975 genotypes in affected and unaffected individuals over generations is given in Supplementary Information S11. The number of individuals carrying the TT does not significantly change over generations in either group. None of the PRS showed a significant difference when comparing PRS for rs1862975 genotypes in affected and unaffected individuals (Supplementary Information S12).

## DISCUSSION

The current study is one of the first the first to probe patterns of common genetic variation within a traditional pedigree design. While increased polygenic scores in patients as compared to unaffected family members have been demonstrated recently (Boies et al. 2017), we aimed to illustrate the possibilities of this approach by investigating apparent assortative mating and anticipation in a large multigenerational pedigree affected with mood disorders through polygenic risk scores for SCZ (Ripke et al. 2014), MDD (PGC, Wray, and Sullivan 2017) and BPD (Stahl et al. 2017) and thereby improve mechanistic understanding of common genetic risk for psychiatric disorders.

Highlighting the possibilities of PRS applications within a family context, we set out to utilize patterns of common variation to illuminate phenomena within the family that are out of reach from traditional case/control studies. Assortative mating is one of the features in this family, where many married-in individuals are more affected with a mood disorder than the general population. As opposed to the family members, the married-in individuals were more often affected with (r)MDD instead of BP. Non-random mating patterns have been reported in the population regarding body type, socio-economic factors and psychiatric traits (Plomin, Krapohl, and O’Reilly 2016; Nordsletten et al. 2016). The BBF provides a unique opportunity to look at the genetic correlation between spouse pairs and the contribution of the offspring of married-in individuals to overall psychiatric morbidity. A recent study has found genetic evidence for assortative mating when studying BMI and height in spouse pairs (Robinson et al. 2017). In the BBF; the affected married-in individuals have a higher, though non-significant, polygenic score than affected or unaffected family members but it appears that we observe significant consequences of this in that the offspring of an affected married-in parent collectively show significantly increased SCZ:PRS and BPD:PRS.

A contribution of the married-in parents to a genetic driven anticipation in age of onset is supported by the increase in SCZ:PRS over generations, although our cross sectional study dataset was less well powered to find an association with age at onset within affected family members. We did observe a trend for association between age at onset and PRS in the youngest generation in this study but not when combining sample across generations. Age at onset can be considered a proxy for severity (Schulze et al. 2002; Schürhoff et al. 2000) and has been previously associated with genetic risk in MDD (Power et al. 2012; Power et al. 2016). However, this variable needs to be interpreted with caution, especially when analyzing patterns over time since it is dependent on context and memory (Alda et al. 2000). Ascertainment bias can be a confounding factor in studies of psychiatric traits, with older generations having less access to psychiatric care and possibly misremembering the onset or nature of their first episode.

Finally, we explored the balance of common and rare risk variation through combining our current PRS results with previously performed linkage analyses. We did not find a decrease in potential rare risk allele genotypes over generations contrasting the increase in SCZ:PRS, and PRS profiles for individuals carrying rare risk genotypes are not significantly different. This indicates that these factors separately confer independent disease risk. Recognizing the limitations in sample size of our pedigree and therefore the power to draw statistically robust conclusions, especially in the combined linkage and PRS analyses, our point is to emphasize the unique nature of the family and we encourage replication in similar pedigrees when available to fully utilize the potential of PRS in this setting.

In conclusion, our study is an exploration of PRS as a tool for investigating patterns of common genetic risk in a traditional pedigree context. The SCZ and BPD scores appear best suited in our data for teasing apart patterns of assortative mating and anticipation, whereby increased polygenic risk for psychiatric disorders is contributed by affected individuals who married into the family, adding to the already present rare risk variation passed on by the early generations (Diniz et al. 2017).

## ACKNOWLEDGEMENTS

The authors would like to thank the family members for their enthusiastic participation. We thank our ethics consultant Prof. Barbara Prainsack for insightful discussions. This paper represents independent research part-funded by FAPESP (2014/50830-2; 2010/08968-6), the Marie Curie International Research Staff Exchange (FP7-PEOPLE-2011-IRSES/295192), and the National Institute for Health Research (NIHR) Biomedical Research Centre at South London and Maudsley NHS Foundation Trust and King’s College London. SDJ is funded by the European Union’s Horizon 2020 research and innovation programme under Marie Skłodowska-Curie grant IF 658195. The views expressed are those of the authors and not necessarily those of the EU, the NHS, the NIHR or the Department of Health. GB has been a consultant in preclinical genomics and has received grant funding from Eli Lilly ltd within the last 3 years. AG has participated in advisory boards for Janssen-Cilag and Daiichi-Sankyo. All other authors report no conflict of interest.

## SUPPLEMENTARY INFORMATION

**Supplementary Information S1:** Full list of Consortium members of the Major Depressive Disorder and Bipolar Disorder working groups of the Psychiatric Genomics Consortium.

**Supplementary Information S2:** Scatterplots of principal components calculated in the Brazilian Bipolar Family members and Brazilian population controls with the Hapmap3 populations.

**Supplementary Information S3:** Scatterplots of principal components calculated in the Brazilian Bipolar Family members and Brazilian population controls coloured by subfamily.

**Supplementary Information S4:** Scatterplots of principal components coloured byy Brazilian Bipolar Family members, married-ins and Brazilian population controls.

**Supplementary Information S5:** Number of children contributed per spouse pair to each offspring category.

**Supplementary Information S6:** List of tests performed.

**Supplementary Information S7:** Violin plots of PRS in diagnostic categories.

**Supplementary Information S8:** Demographics of offspring from different spouse pair categories.

**Supplementary Information S9:** Violin plot of age at onset over generations.

**Supplementary Information S10:** Scatterplots of PRS and age at onset for affected family members.

**Supplementary Information S11:** Stacked barplots showing the genotype distributions of the risk SNP over generations for affected and unaffected family members.

**Supplementary Information S12:** Violin plots of PRS for groups carrying different risk SNP genotypes.

## REFERENCES

Abecasis, Gonçalo R, Stacey S Cherny, William O Cookson, and Lon R Cardon. 2002. “Merlin–Rapid Analysis of Dense Genetic Maps Using Sparse Gene Flow Trees.” Nature Genetics 30 (1): 97–101. doi:10.1038/ng786.

Ahn, K, S S An, Y Y Shugart, and J L Rapoport. 2016. “Common Polygenic Variation and Risk for Childhood-Onset Schizophrenia.” Molecular Psychiatry 21 (1): 94–96. doi:10.1038/mp.2014.158.

Alda, M, P Grof, L Ravindran, P Cavazzoni, A Duffy, E Grof, P Zvolský, and J Wilson. 2000. “Anticipation in Bipolar Affective Disorder: Is Age at Onset a Valid Criterion?” American Journal of Medical Genetics 96 (6): 804–7. http://www.ncbi.nlm.nih.gov/pubmed/11121186.

Boies, Sébastien, Chantal Mérette, Thomas Paccalet, Michel Maziade, and Alexandre Bureau. 2017. “Polygenic Risk Scores Distinguish Patients from Non-Affected Adult Relatives and from Normal Controls in Schizophrenia and Bipolar Disorder Multi-Affected Kindreds.” American Journal of Medical Genetics Part B: Neuropsychiatric Genetics, November. doi:10.1002/ajmg.b.32614.

Brasil, Heloisa H A, and Isabel A Bordin. 2010. “Convergent Validity of K-SADS-PL by Comparison with CBCL in a Portuguese Speaking Outpatient Population.” BMC Psychiatry 10 (January): 83. doi:10.1186/1471-244X-10-83.

Butler, D G, B R Cullis, A R Gilmour, and B J Gogel. 2002. “ASReml-R Reference Manual 3.” Tech. Report, Dep. Prim. Ind. Queensl.

Chang, Christopher C, Carson C Chow, Laurent CAM Tellier, Shashaank Vattikuti, Shaun M Purcell, James J Lee, S Purcell, et al. 2015. “Second-Generation PLINK: Rising to the Challenge of Larger and Richer Datasets.” GigaScience 4 (1). BioMed Central: 7. doi:10.1186/s13742-015-0047-8.

Cross-Disorder Group of the Psychiatric Genomics Consortium. 2013. “Identification of Risk Loci with Shared Effects on Five Major Psychiatric Disorders: A Genome-Wide Analysis.” Lancet 381 (9875): 1371–79. doi:10.1016/S0140-6736(12)62129-1.

Del-Ben, C M, C R Rodrigues, and A W Zuardi. 1996. “Reliability of the Portuguese Version of the Structured Clinical Interview for DSM-III-R (SCID) in a Brazilian Sample of Psychiatric Outpatients.” Brazilian Journal of Medical and Biological Research = Revista Brasileira de Pesquisas Médicas E Biológicas / Sociedade Brasileira de Biofísica … [et Al.] 29 (12): 1675–82. http://www.ncbi.nlm.nih.gov/pubmed/9222432.

Diniz, Mateus Jose Abdalla, Andiara Calado Saloma Rodrigues, Ary Gadelha, Shaza Alsabban, Camila Guindalini, Jose Paya-Cano, Simone de Jong, Peter McGuffin, Rodrigo Affonseca Bressan, and Gerome Breen. 2017. “Whole Genome Linkage Analysis in a Large Brazilian Multigenerational Family Reveals Distinct Linkage Signals for Bipolar Disorder and Depression.” bioRxiv 106260.

Euesden, Jack, Cathryn M Lewis, and Paul F O’Reilly. 2014. “PRSice: Polygenic Risk Score Software.” Bioinformatics (Oxford, England), December. doi:10.1093/bioinformatics/btu848.

Kenward, M G, and J H Roger. 1997. “Small Sample Inference for Fixed Effects from Restricted Maximum Likelihood.” Biometrics 53 (3): 983–97. http://www.ncbi.nlm.nih.gov/pubmed/9333350.

Lee, S Hong, Stephan Ripke, Benjamin M Neale, Stephen V Faraone, Shaun M Purcell, Roy H Perlis, Bryan J Mowry, et al. 2013. “Genetic Relationship between Five Psychiatric Disorders Estimated from Genome-Wide SNPs.” Nature Genetics 45 (9). Nature Publishing Group, a division of Macmillan Publishers Limited. All Rights Reserved.: 984–94. doi:10.1038/ng.2711.

Lichtenstein, Paul, Benjamin H Yip, Camilla Björk, Yudi Pawitan, Tyrone D Cannon, Patrick F Sullivan, and Christina M Hultman. 2009. “Common Genetic Determinants of Schizophrenia and Bipolar Disorder in Swedish Families: A Population-Based Study.” Lancet 373 (9659): 234–39. doi:10.1016/S0140-6736(09)60072-6.

Merikangas, K R, and D G Spiker. 1982. “Assortative Mating among in-Patients with Primary Affective Disorder.” Psychological Medicine 12 (4): 753–64. http://www.ncbi.nlm.nih.gov/pubmed/7156249.

Nakagawa, Shinichi, and Holger Schielzeth. 2013. “A General and Simple Method for Obtaining R 2 from Generalized Linear Mixed-Effects Models.” Edited by Robert B. O’Hara. Methods in Ecology and Evolution 4 (2): 133–42. doi:10.1111/j.2041-210x.2012.00261.x.

Nordsletten, Ashley E, Henrik Larsson, James J Crowley, Catarina Almqvist, Paul Lichtenstein, and David Mataix-Cols. 2016. “Patterns of Nonrandom Mating Within and Across 11 Major Psychiatric Disorders.” JAMA Psychiatry 73 (4): 354–61. doi:10.1001/jamapsychiatry.2015.3192.

Noto, Cristiano, Vanessa Kiyomi Ota, Marcos Leite Santoro, Bruno B Ortiz, Lucas B Rizzo, Cinthia Hiroko Higuchi, Quirino Cordeiro, et al. 2015. “Effects of Depression on the Cytokine Profile in Drug Naïve First-Episode Psychosis.” Schizophrenia Research 164 (1-3): 53–58. doi:10.1016/j.schres.2015.01.026.

O’Donovan, Michael, Ian Jones, and Nick Craddock. 2003. “Anticipation and Repeat Expansion in Bipolar Disorder.” American Journal of Medical Genetics. Part C, Seminars in Medical Genetics 123C (1): 10–17. doi:10.1002/ajmg.c.20009.

Patterson, Nick, Alkes L Price, and David Reich. 2006. “Population Structure and Eigenanalysis.” Edited by David B Allison. PLoS Genetics 2 (12). Public Library of Science: e190. doi:10.1371/journal.pgen.0020190.

PGC, - Major Depressive Disorder Working Group of the, Naomi R. Wray, and Patrick F Sullivan. 2017. “Genome-Wide Association Analyses Identify 44 Risk Variants and Refine the Genetic Architecture of Major Depression.” bioRxiv, July. Cold Spring Harbor Laboratory, 167577. doi:10.1101/167577.

Plomin, Robert, Eva Krapohl, and Paul F. O’Reilly. 2016. “Assortative Mating—A Missing Piece in the Jigsaw of Psychiatric Genetics.” JAMA Psychiatry 73 (4): 323–24. doi:10.1001/jamapsychiatry.2015.3204.

Power, Robert A., Katherine E. Tansey, Henriette Nørmølle Buttenschøn, Sarah Cohen-Woods, Tim Bigdeli, Lynsey S. Hall, Zoltán Kutalik, et al. 2016. “Genome-Wide Association for Major Depression through Age at Onset Stratification.” Biological Psychiatry, May. doi:10.1016/j.biopsych.2016.05.010.

Power, Robert A, Robert Keers, Mandy Y Ng, Amy W Butler, Rudolf Uher, Sarah Cohen-Woods, Marcus Ising, et al. 2012. “Dissecting the Genetic Heterogeneity of Depression through Age at Onset.” American Journal of Medical Genetics. Part B, Neuropsychiatric Genetics: The Official Publication of the International Society of Psychiatric Genetics 159B (7): 859–68. doi:10.1002/ajmg.b.32093.

Purcell, S M, N R Wray, J L Stone, P M Visscher, M C O’Donovan, P F Sullivan, and P Sklar. 2009. “Common Polygenic Variation Contributes to Risk of Schizophrenia and Bipolar Disorder.” Nature 460 (7256): 748–52. http://www.ncbi.nlm.nih.gov/entrez/query.fcgi?cmd=Retrieve&db=PubMed&dopt=Citation&list_uids=19571811.

Purcell, S, B Neale, K Todd-Brown, L Thomas, M A Ferreira, D Bender, J Maller, et al. 2007. “PLINK: A Tool Set for Whole-Genome Association and Population-Based Linkage Analyses.” Am J Hum Genet 81 (3): 559–75. http://www.ncbi.nlm.nih.gov/entrez/query.fcgi?cmd=Retrieve&db=PubMed&dopt=Citation&list_uids=17701901.

Purcell, Shaun M, Naomi R Wray, Jennifer L Stone, Peter M Visscher, Michael C O’Donovan, Patrick F Sullivan, and Pamela Sklar. 2009. “Common Polygenic Variation Contributes to Risk of Schizophrenia and Bipolar Disorder.” Nature 460 (7256): 748–52. doi:10.1038/nature08185.

Rioux, John D., Mark J. Daly, Mark S. Silverberg, Kerstin Lindblad, Hillary Steinhart, Zane Cohen, Terrye Delmonte, et al. 2001. “Genetic Variation in the 5q31 Cytokine Gene Cluster Confers Susceptibility to Crohn Disease.” Nature Genetics 29 (2): 223–28. doi:10.1038/ng1001-223.

Ripke, Stephan, Benjamin M. Neale, Aiden Corvin, James T. R. Walters, Kai-How Farh, Peter A. Holmans, Phil Lee, et al. 2014. “Biological Insights from 108 Schizophrenia-Associated Genetic Loci.” Nature 511 (7510): 421–27. doi:10.1038/nature13595.

Robinson, Matthew R., Aaron Kleinman, Mariaelisa Graff, Anna A. E. Vinkhuyzen, David Couper, Michael B. Miller, Wouter J. Peyrot, et al. 2017. “Genetic Evidence of Assortative Mating in Humans.” Nature Human Behaviour 1 (1). Nature Publishing Group: 16. doi:10.1038/s41562-016-0016.

Schulze, Thomas G, Daniel J Müller, Harald Krauss, Magdalena Gross, Heiner Fangerau-Lefèvre, Franciska Illes, Stephanie Ohlraun, et al. 2002. “Further Evidence for Age of Onset Being an Indicator for Severity in Bipolar Disorder.” Journal of Affective Disorders 68 (2-3): 343–45. http://www.ncbi.nlm.nih.gov/pubmed/12063163.

Schürhoff, F, F Bellivier, R Jouvent, M C Mouren-Siméoni, M Bouvard, J F Allilaire, and M Leboyer. 2000. “Early and Late Onset Bipolar Disorders: Two Different Forms of Manic-Depressive Illness?” Journal of Affective Disorders 58 (3): 215–21. http://www.ncbi.nlm.nih.gov/pubmed/10802130.

Silva, Marcus T, Tais F Galvao, Silvia S Martins, and Mauricio G Pereira. 2014. “Prevalence of Depression Morbidity among Brazilian Adults: A Systematic Review and Meta-Analysis.” Revista Brasileira de Psiquiatria (Sao Paulo, Brazil: 1999) 36 (3): 262–70. http://www.ncbi.nlm.nih.gov/pubmed/25119639.

Speed, Doug, Gibran Hemani, Michael R Johnson, and David J Balding. 2012. “Improved Heritability Estimation from Genome-Wide SNPs.” American Journal of Human Genetics 91 (6). Elsevier: 1011–21. doi:10.1016/j.ajhg.2012.10.010.

Stahl, Eli, Andreas Forstner, Andrew McQuillin, Stephan Ripke, Bipolar Disorder Working Group of the PGC, Roel Ophoff, Laura Scott, et al. 2017. “Genomewide Association Study Identifies 30 Loci Associated with Bipolar Disorder.” bioRxiv, August. Cold Spring Harbor Laboratory, 173062. doi:10.1101/173062.

Sullivan, Patrick F. 2010. “The Psychiatric GWAS Consortium: Big Science Comes to Psychiatry.” Neuron 68 (2): 182–86. doi:10.1016/j.neuron.2010.10.003.

Berk, M., S. Dodd, P. Callaly, L. Berk, P. Fitzgerald, A.R. de Castella, S. Filia, et al. 2007. “History of Illness prior to a Diagnosis of Bipolar Disorder or Schizoaffective Disorder.” Journal of Affective Disorders 103 (1-3): 181–86. doi:10.1016/j jad.2007.01.027.

Lewis, Cathryn M. 2015. “Dissecting the Genetic Contribution to Depression: Progress at Last.” In World Congress of Psychiatric Genetics.

Plomin, Robert, Eva Krapohl, and Paul F. O’Reilly. 2016. “Assortative Mating–A Missing Piece in the Jigsaw of Psychiatric Genetics.” JAMA Psychiatry 73 (4): 323–24. doi:10.1001/jamapsychiatry.2015.3204.

Del-Ben, C M, C R Rodrigues, and A W Zuardi. 1996. “Reliability of the Portuguese Version of the Structured Clinical Interview for DSM-III-R (SCID) in a Brazilian Sample of Psychiatric Outpatients.” Brazilian Journal of Medical and Biological Research = Revista Brasileira de Pesquisas Médicas E Biológicas / Sociedade Brasileira de Biofísica … [et al.] 29 (12): 1675–82. http://www.ncbi.nlm.nih.gov/pubmed/9222432.

Sklar, Pamela. 2015. “Genome-Wide Association Study Using Psychchip in a Cohort of >13,000 New Bipolar Disorder Cases.” In World Congress of Psychiatric Genetics. doi:10.1016/j.euroneuro.2015.09.009.

